# A Kernel Method for Dissecting Genetic Signals in Tests of High-Dimensional Phenotypes

**DOI:** 10.1101/2021.07.29.454336

**Authors:** Claudia Solis-Lemus, Aaron M. Holleman, Andrei Todor, Bekh Bradley, Kerry J. Ressler, Debashis Ghosh, Michael P. Epstein

**Affiliations:** Department of Human Genetics, Emory University, Atlanta, GA; Department of Epidemiology, Emory University, Atlanta, GA; Department of Biostatistics and Informatics, University of Colorado, Aurora, CO; Department of Psychiatry and Behavioral Sciences, Emory University, Atlanta, GA; Clinical Psychologist, Mental Health Service Line, Department of Veterans Affairs Medical Center, Atlanta, GA; Department of Psychiatry, McLean Hospital, Harvard Medical School, Belmont, MA

**Author notes:** Address for correspondence: Dr. Michael Epstein, Department of Human Genetics, Emory University School of Medicine, Atlanta, GA, 30030.

**Keywords:** pleiotropy, association, high dimension, complex human traits

## Abstract

Genomewide association studies increasingly employ multivariate tests of multiple correlated phenotypes to exploit likely pleiotropy to improve power. Typical multivariate methods produce a global p-value of association between a variant (or set of variants) and multiple phenotypes. When the global test is significant, subsequent interest then focuses on dissecting the signal and, in particular, delineating the set of phenotypes where the genetic variant(s) have a direct effect from the remaining phenotypes where the genetic variant(s) possess either indirect or no effect. While existing techniques like mediation models can be utilized for this purpose, they generally cannot handle high-dimensional phenotypic and genotypic data. To assist in filling this important gap, we propose a modification of a kernel distance-covariance framework for gene mapping of multiple variants with multiple phenotypes to test instead whether the association between the variants and a group of phenotypes is driven through a direct association with just a subset of the phenotypes. We use simulated data to show that our new method controls for type I error and is powerful to detect a variety of models demonstrating different patterns of direct and indirect effects. We further illustrate our method using GWAS data from the Grady Trauma Project and show that an existing signal between genetic variants in the ZHX2 gene and 21 items within the Beck Depression Inventory appears to be due to a direct effect of these variants on only 3 of these items. Our approach scales to genomewide analysis, and is applicable to high-dimensional correlated phenotypes.

## 1 Introduction

Genetic analysis of multiple correlated phenotypes simultaneously is a popular strategy for gene mapping owing to the observation that many trait-influencing variants can influence multiple distinct phenotypes through pleiotropy (see Solovieff et al. (2013), Galesloot et al. (2014), Chung et al. (2014)). The difficulty in most multivariate tests lies in the fact that they are omnibus tests, and thus, these tests do not provide insight on which of the phenotypes in the set are driving the multivariate association through a direct association with the genotypes (direct effect), which phenotypes are only associated with the genotypes due to mediation (indirect effect), and which phenotypes are not associated with the genotypes. Such information would be valuable to obtain to gain important understanding of underlying biological processes.

A general strategy to distinguish direct effects from indirect effects in studies of association between exposures and outcomes of interest involves mediation analyses. In genetic studies, researchers apply mediation analyses to assess whether a cross-phenotype effect is due to biological pleiotropy or mediation pleiotropy. For example, suppose we find a significant cross-phenotype association between a SNP and two phenotypes (P1 and P2). Biological pleiotropy states that the SNP has direct effects on both P1 and P2. Mediation, on the other hand, considers the possibility that perhaps P2 lies in the pathway between the SNP and P1 such that at least part of the association between SNP and P1 is indirect due to each having a direct relationship with P2. Such mediation analyses are increasingly common in genetic studies of complex traits (Huang et al., 2014; VanderWeele et al., 2012; Gabrielsen et al., 2013).

Existing methods for mediation analysis are not tailored to handle situations where the outcome variable, mediator variable, and genetic variable are of possibly high dimension. Thus, current methods are not applicable in a situation that involves testing whether a group of rare variants is only directly associated with a (possibly high dimensional) subset of the original collection of phenotypes. To fill this important gap, we propose a test for dissecting those phenotypes driving a multivariate association signal through direct associations with a genotype or set of genotypes in a gene. Our idea builds on GAMuT (Broadaway et al., 2016), a powerful method for cross-phenotype testing of both common and rare variants in a gene using a kernel distance-covariance (KDC) framework (Gretton et al., 2008; Székely et al., 2007; Székely and Rizzo, 2009; Hua and Ghosh, 2015). Our new method (“GAMuT-Dissect”) identifies whether an observed association between a variant (or a group of variants within a gene) and possibly high-dimensional phenotype data is driven by direct association with a specific subset of these phenotypes (also possibly of high dimension). The method can handle both categorical and continuous data and further can adjust for covariates. Our method also yields analytic p-values without the need of permutation or resampling, thereby facilitating large-scale analysis.

The organization of the manuscript is as follows: in section 2.1, we describe the methods for the pleiotropic analysis of high-dimensional phenotypes (from Broadaway et al. (2016)). In section 2.2, we describe the extension of GAMuT to enable dissection of direct effects among high-dimensional phenotypes. In section 3, we present simulation results demonstrating the feasibility of the approach. We further illustrate the method by applying the technique to genetic and high-dimensional phenotypic data from the Grady Trauma Project dataset to dissect a previously detected association between the gene ZHX2 and the 21-item Beck Depression Inventory. Using our method, we show that the gene may have direct association with only 3 of the 21 items that compose this phenotype. Finally, in section 4, we elaborate on some limitations of our methodology and future areas of improvement.

## 2 Materials and Methods

### 2.1 GAMuT for pleiotropic analysis of high-dimensional phenotypes

GAMuT (Broadaway et al., 2016) studies the association between high-dimensional phenotypes and highdimensional genotypes, through a nonparametric test of independence between two sets of multivariate variables. We assume a sample of *N* subjects that are genotyped for *V* genetic variants in a target gene or region. We further assume these subjects are measured for *L* phenotypes. For subject *j* (*j* = 1,…, *N*), let **Y**_*j*_. = (*Y*_*j*,1_,*Y*_*j*,2_,…, *Y*_*j*,*L*_) be the *L*-dimensional vector of phenotypes, so that 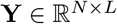 is the full matrix of phenotypes. Similarly, let *G_j_*. = (*G*_*j*,1_, *G*_*j*,2_,…, *G*_*j*,*V*_) be the genotypes of subject *j* at *V* variants in the gene or region of interest. Note that *G*_*j*,*v*_ represents the number of copies of the minor allele that the subject possesses at the *v^th^* variant. Thus, the matrix of genotypes for the sample is denoted 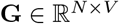.

GAMuT tests for independence between **Y** (the *N* × *L* matrix of multivariate phenotypes) and **G** (the *N* × *V* matrix of multivariate genotypes) by constructing an *N* × *N* phenotypic-similarity matrix **Q**, and an *N* × *N* genotypic-similarity matrix **X**. See Broadaway et al. (2016) for more details on how to model pairwise similarity or dissimilarity.

After constructing the similarity matrices **Q** and **X**, we center them as **Q**_*c*_ = **HQH** and **X**_*c*_ = **HXH**, where **H** = (**I** – **11**^*T*^/*N*) is a centering matrix (**HH** = **H**), 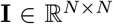 is an identity matrix, and 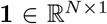 is a vector of ones. With the centered similarity matrices (**Q**_*c*_, **X**_*c*_), we construct the GAMuT test statistic as

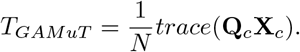

Under the null hypothesis that the two matrices are independent, *T_GAMuT_* follows the asymptotic distribution of a weighted sum of independent and identically distributed 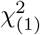 variables (Broadaway et al., 2016). We then use Davies’ method (Davies, 1980) to analytically calculate the p-value of *T_GAMuT_*.

Our framework for determining those phenotypes driving a multivariate association signal through direct associations with a set of genotypes involves two separate hypothesis tests: 1) standard GAMuT test to identify if there is association between *G* and *Y* (as described in this section), and 2) GAMuT-Dissect to identify if there is association between *P*_1_ (a subset of phenotypes) and *G*, *after removing the effect of the potential mediator P*_2_ (which will be described in the next section).

### 2.2 GAMuT-Dissect for determining subset of high-dimensional phenotypes with direct genetic effects

Let 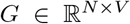 be a genotype matrix of *N* unrelated individuals and *V* genetic variants. Let 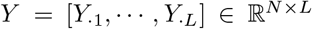 be a phenotype matrix containing *L* phenotype vector columns 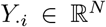. Let *Y* = [*P*_1_, *P*_2_] be a partition of the phenotype matrix with 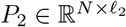 as the potential mediator between *G* and 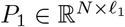, with *ℓ*_1_ + *ℓ*_2_ = *L* (see Figure 1). Without loss of generality, we can assume that *Y_·i_* ∈ *P*_1_ for *i* = 1,…, *ℓ*_1_ and *Y_·i_* ∈ *P*_2_ for *i* = *ℓ*_1_ + 1,…, *L*. Let 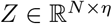 represent the remaining shared (e.g. polygenic) effect that affects *P*_1_ and *P*_2_ simultaneously (see Figure 1). This effect drives the non-causal association between *P*_1_ and *P*_2_. Without loss of generality, we assume here that the shared effect is one-dimensional, and thus, *η* =1.

**Figure 1:**
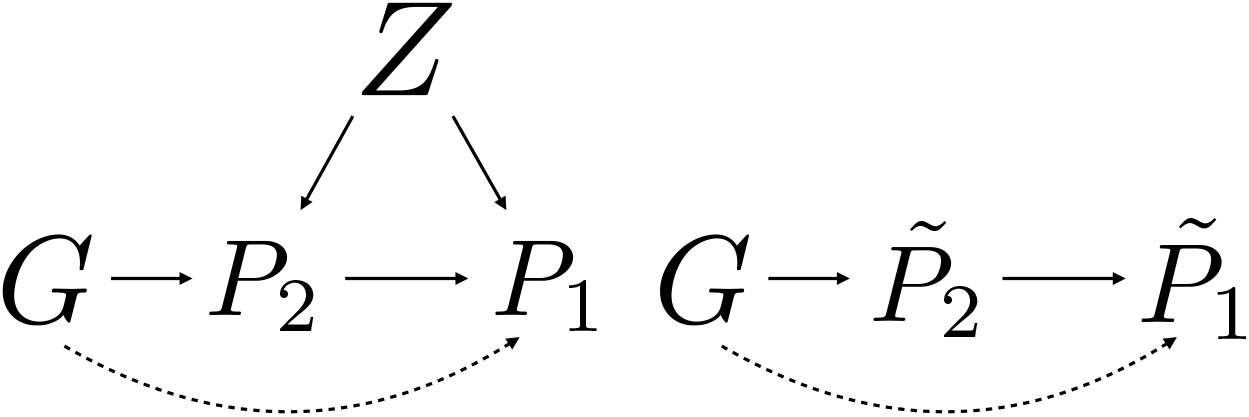
**Left:** *Z* is a latent variable that represents the polygenic effect affecting both *P*_1_ and *P*_2_. If the dashed line is absent, the genotype *G* is only directly associated with *P*_2_, so any association between *G* and *P*_1_ is due to mediation or correlation. **Right:** 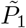 and 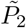 represent the residuals after regressing out the latent variable *Z*.

GAMuT-Dissect tests the null hypothesis that a direct association between *G* and *P*_2_ fully explains the previously observed association between *G* and *Y* (no dashed line in Figure 1), after removing the effect of *Z* if we believe such an effect to exist. If *Z* is believed to exist and is measurable, we can regress out this effect from each phenotype under consideration (see Algorithm 1). If *Z* is instead assumed to be random, one could possibly eigendecompose the resulting covariance matrix and multiply the phenotypes by the eigenvector matrix (Zhou and Stephens, 2014). We detail this idea more in Discussion.

Under the null hypothesis (*H*_0_: 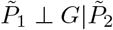), 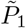 should not be associated with *G* after conditioning on 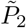, suggesting that 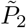 explains the previously observed association between *G* and *Y*. The alternative hypothesis is that there is association between *G* and 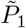 after conditioning on 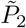, indicating that 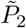 either 1) explains some but not all of the previously observed association between *G* and *Y* or 2) 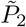 does not contribute to the association between *G* and *Y*.

Note that in the algorithm (Algorithm 1), we first create the kernel matrix for the phenotypes and genotypes based on Broadaway et al. (2016). A kernel matrix is a matrix obtained by applying a pairwise kernel function on the phenotype or genotype matrix. Among the kernel functions, we can highlight the linear kernel or projection kernel, but see Broadaway et al. (2016) and references therein for more details.

Finally, we note that this method can also be applied to chromosome X variant sets by following the genotype coding in Ma et al. (2015).

#### Algorithm 1: GAMuT-Dissect

**Input:** Genotype matrix *G*, Phenotype matrix *Y* = [*P*_1_, *P*_2_], where *P*_2_ is the potential mediator, *Z* polygenic effects on *Y*

- For every column *Y_·i_* in *Y*, let *Ỹ_i_* be the residuals of the linear regression *Y_·i_* ~ *Z*. If there are covariates in the dataset, *Y_·i_* represents the residuals after regressing out the covariates.

- Let 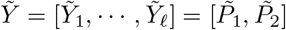 be the new matrix of phenotypes, after the effect of *Z* has been removed

- Let *K*_1_ ∈ **R**^*N*×*N*^ represent the (linear or projection) kernel matrix of 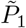, and *K_G_* ∈ **R**^*N*×*N*^ the (linear or projection) kernel matrix of the genotype matrix *G*. For each column *i* in *K*_1_ and *K_G_*, we will fit a linear regression model on 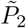 and extract the residuals. That is, for column *i*, let 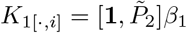 denote the linear regression model, where **1** ∈ **R**^*N*^, *β*_1_ ∈ **R**^*ℓ*_2_+1^, the fitted values are given by 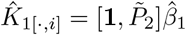, and residuals are given by 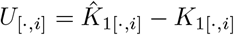 and stored in the *i^th^* column of the matrix *U* ∈ **R**^*N*×*N*^. Similarly, for column *i*, let 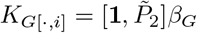 denote the linear regression model, where **1** ∈ **R**^*N*^, *β_G_* ∈ **R**^*ℓ*_2_+1^, the fitted values are given by 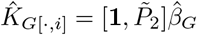, and residuals are given by 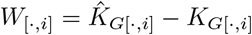 and stored in the *i^th^* column of the matrix *W* ∈ **R**^*N*×*N*^.

- Perform a standard GAMuT test on the matrices *W* and *U*

**Output:** P-value of the hypothesis test of independence between *W* and *U*.

### 2.3 Adjusting for covariates

It is important for pleiotropic tests to adjust for important covariates, such as principal components of ancestry, to avoid potential confounding. The adjustment for covariates in GAMuT-Dissect is handled in a similar manner to that in the standard GAMuT (Broadaway et al., 2016). Before applying GAMuT, we can control for confounders by regressing each phenotype on covariates of interest and then using the residuals to form the phenotypic similarity matrix. We note that even though such residualization is not usual for the case of binary phenotypes, some studies have shown the validity of this procedure in genetic association studies (Price et al., 2006; Kang et al., 2010, Wei and Lu (2017)). Finally, GAMuT-Dissect can also handle potential cryptic relatedness in a similar fashion to the standard GAMuT by eigendecomposing the sample relatedness matrix and multiplying the phenotypes and genotypes by the eigenvector matrix (Zhou and Stephens, 2014).

### 2.4 Simulations

We conducted simulations to show that GAMuT-Dissect properly preserves type I error and to further assess power under different models of direct, indirect, and no effects of genotype. Under the simulation setup in Figure 2 (left), we used R to simulate a region of 500 independent genetic variants (assuming 1% to be causal) with varying MAF between 0.01 and 0.05 (see the SM for MAF equal to 0.25). We simulated the polygenic effects *Z* as independent normally distributed with mean 0 and variance 1. We simulated 4 continuous phenotypes: (*Y*_1_, *Y*_2_, *Y*_3_, *Y*_4_), with *Y*_1_ and *Y*_2_ associated with *G* through two different effect sizes: *α*_1_ for *Y*_1_ and *α*_2_ for *Y*_2_. All 4 phenotypes are correlated through the latent scalar variable *Z* which represents the remaining polygenic effects not captured by *G*, with a specific effect size *β* (assumed *β_i_* = *β* for all *i* = 1, 2, 3, 4). This remaining polygenic effect is assumed known here. We varied *β* as 0, 0.25, 0.5, 1.0. For the case of rare variants, the effect sizes (*α_i_*) were multiplied by | log_10_(*MAF*)| so that the effect size of a given causal variant is inversely proportional to its MAF. For an individual *k*, the vector (*Y*_*k*1_, *Y*_*k*2_, *Y*_*k*3_, *Y*_*k*4_) is drawn from a multivariate normal with mean *α_i_G_k_* + *βZ_k_* for *Y_i_* (*i* =1, 2), and mean *βZ_k_* for *Y*_3_, *Y*_4_. The covariance matrix given by 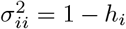 and 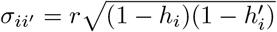 where *r* was randomly sampled from a uniform distribution on (0.3, 0.5) (medium-sized correlation) and with *h_i_* = ∑2*α_i_MAF* (1 – *MAF*) where the sum is over the causal genetic variants (*i* = 1, 2, 3, 4). For *i* = 3, 4, *alpha_i_* is set as 0 in the definition of *h_j_*.

**Figure 2:**
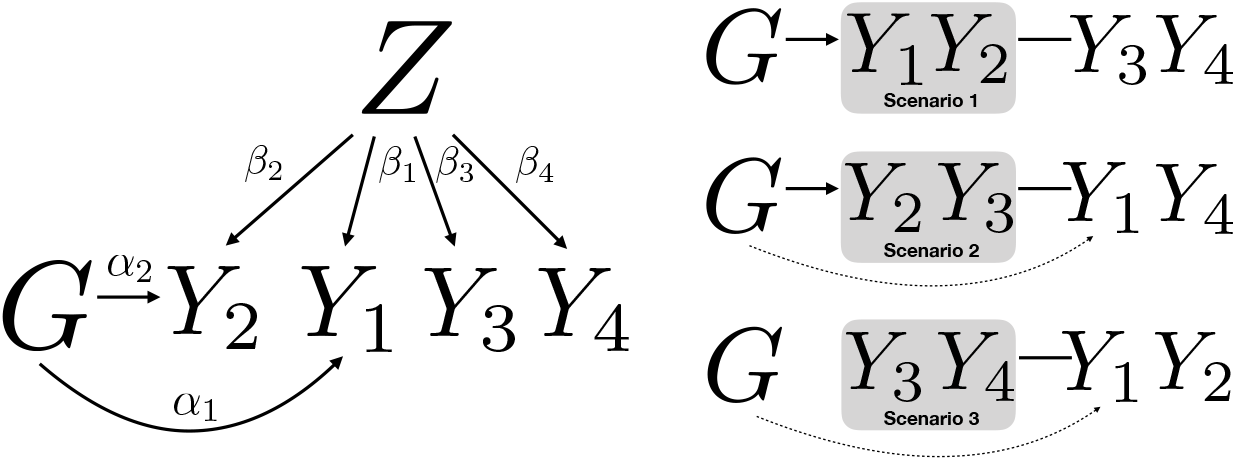
**Left:** Simulation scenario with 4 phenotypes and set of genetic variants *G*. **Right:** Three scenarios tested. Scenario 1: *G* directly associated with intermediate variables only, Scenario 2: *G* directly associated with intermediate and outcome variables, and Scenario 3: *G* directly associated with outcome variables only.

The effect sizes were varied within the range *α_i_* ∈ (0.0, 0.11) for *i* = 1, 2 so that the *R*^2^ was kept between (0, 0.027). The 4 phenotypes were split in two groups: intermediate group and outcome group under three scenarios (Figure 2 right). Scenario 1 models *G* as having direct effects on intermediate variables only. Scenario 2 models *G* as having direct effects on both intermediate and outcome variables. Finally, Scenario 3 assumes *G* has direct effects on outcome variables only.

We used a sample size of 5000 unrelated individuals, and replicated each simulation setting 100 times for power simulations and 1000 for null simulations. The empirical power was estimated by computing the proportion of p-values less than the significance level (*α* = 0.05) out of 100 replicates per scenario.

### 2.5 Analysis of the Grady Trauma Project

The Grady Trauma Project (GTP) studies genetic risk factors associated with psychiatric disorders such as PTSD and depression (Bradley et al., 2008; Ressler et al., 2011) (protocols approved by the IRBs of Emory University School of Medicine and Grady Memorial Hospital). The participants in GTP are predominantly African-American of low socioeconomic status, served by the Grady Hospital in Atlanta, Georgia. GTP staff collected an Oragene salivary sample for DNA extraction. Participants were genotyped on the Illumina HumanOmni1-Quad array for GWAS analyses. After standard GWAS quality control filters, 4,607 African-American subjects remained in the dataset with good quality genotype data.

GTP staff also conducts an extensive verbal interview, with demographic information, history of stressful life events, and several psychological surveys, including the Beck Depression Inventory (BDI). The BDI contains 21 groups of statements that represent diverse symptoms and attitudes associated with depression. Each group includes 4 statements with intensity in a scale of 0 to 3. The 21 groups are in Table 1.

**Table 1:**
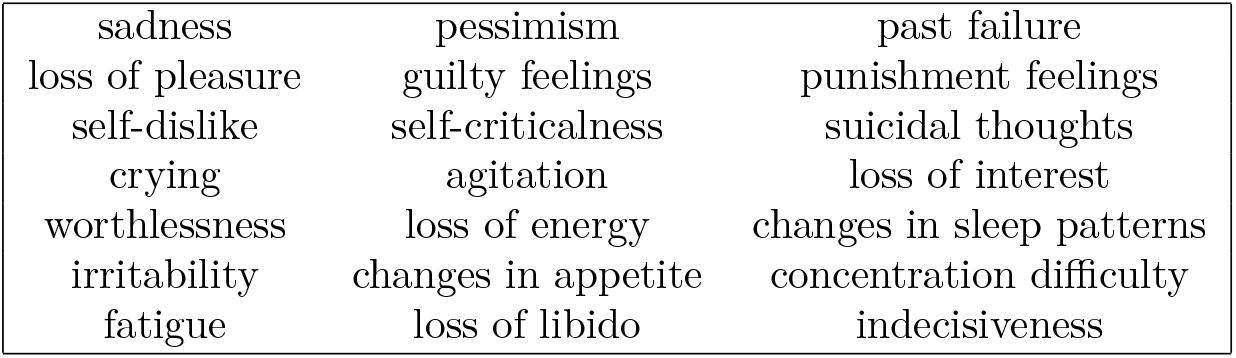
21 Items that Constitute the Beck Depression Inventory (BDI)

The BDI is generally self-administered or self-reported, and is scored by summing the ratings given to each of the 21 items. Summing the responses yields a score ranging from 0-63, with scores higher than 28 being indicative of moderate to severe depression. Subjects who did not report at least one past trauma, subjects with missing BDI scores, or subjects with incomplete covariate data (age, gender, and the top ten principal components to account for ancestry) were also removed, and thus, the final sample contained 3,520 subjects.

Holleman et al. (2019) applied GAMuT to the BDI data using a linear kernel to measure pairwise phenotypic similarity in multivariate symptom scores on 765,580 common genetic variants (MAF > 5%) that fell within 19,609 autosomal genes. Their analyses identified one gene exceeding study-wise significance: ZHX2, on chromosome 8 was strongly associated with the symptoms in the BDI (*P* = 2.73 × 10^-6^). Previous studies have identified a possible link between ZHX2 and autism spectrum disorder (Walker and Scherer, 2013). Here, we used GAMuT-Dissect to explore whether the significant association between ZHX2 and the 21 symptoms within the BDI is driven by direct associations of the variants in this gene with only a select subset of the symptoms.

For the real-life data application, the set of mediators is not known in advance. We propose a sequential approach in which first each individual phenotype is tested as a potential mediator (with the remaining *L* – 1 in *P*_1_). Then, the set of potential mediators is created by including those phenotypes identified as mediators by the GAMuT-Dissect test. In a second stage, pairs of mediators are tested together in *P*_2_ (with the remaining *L* – 2 phenotypes in *P*_1_). This same procedure is followed until the largest possible set of mediators is identified by GAMuT-Dissect. We then adjust for multiple testing (Schaid et al., 2016; Stevens et al., 2017). See the Discussion for more details on the limitations of the sequential approach.

## 3 Results

### 3.1 Type I error simulations (Scenario 1)

Figure 3 shows the quantile-quantile (QQ) plots of 1000 null simulations of Scenario 1 (Figure 2) for our GAMuT-Dissect test when we adjust by the polygenic effect (*Z*) on all the phenotypes (as described in algorithm 1). We do not observe any inflation of p-values, and thus, the type I error for the null hypothesis of Scenario 1 is well-preserved. Thus, our method is properly calibrated for the situation where the genetic variants have direct effects only on the intermediate variables.

**Figure 3:**
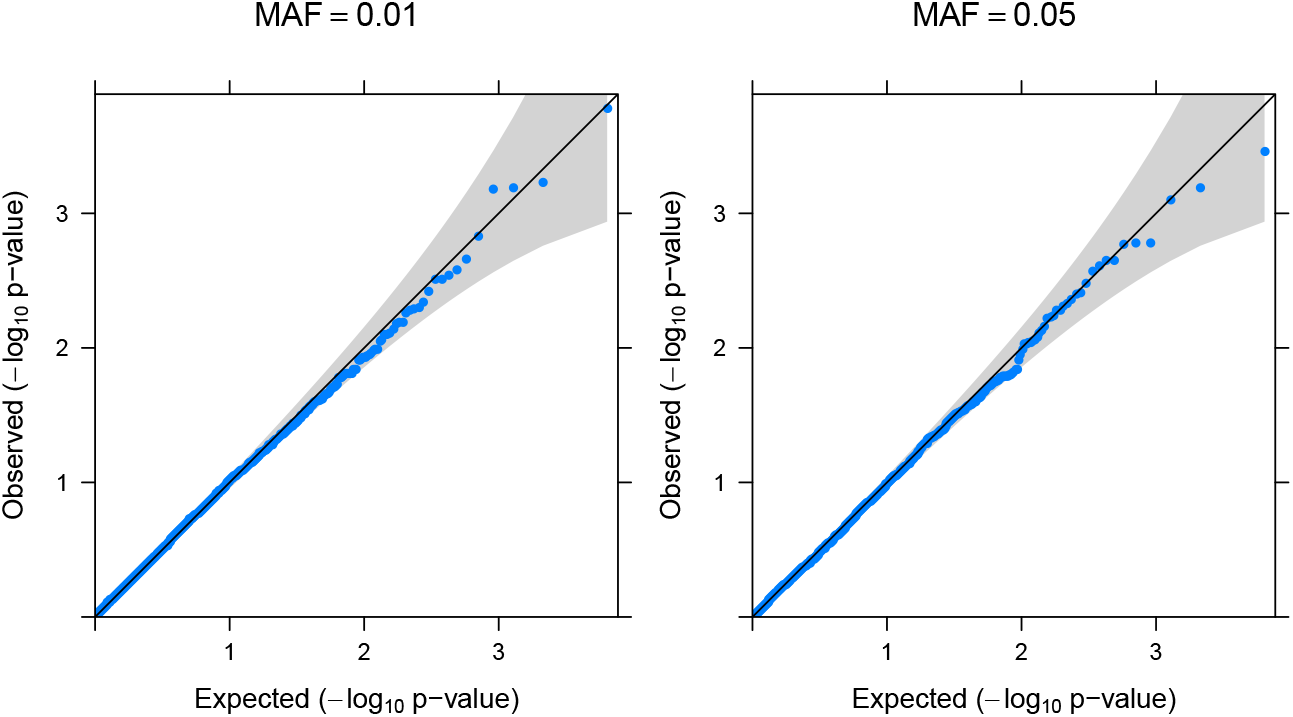
Type I error simulations: Scenario 1. Q-Q plots of p-values for null simulations of scenario 1 for GAMuT-Dissect. Simulated datasets (1000) assumed a window of 500 genetic variants with a given MAF (0.01 or 0.05), assuming 1% to be causal.

### 3.2 Power simulations (Scenario 2)

Here, we show that GAMuT-Dissect has power to detect Scenario 2 where genetic variants have direct effects on both intermediate and outcome variables (see Figure 2 right) after controlling for polygenic effect *Z*.

We present power plots as a function of *R*^2^ corresponding to *α*_1_, because this effect is the one driving the simulation model, with different lines for *R*^2^ corresponding to *α*_2_. Figure 4 shows that GAMuT-Dissect has increasing power as the effect *α*_1_ increases, regardless of the value of *α*_2_. We also show that the parameters chosen for the effect sizes (*α*_1_, *α*_2_) produce datasets with enough power to detect omnibus association with *G* (standard GAMuT in Figure 5).

**Figure 4:**
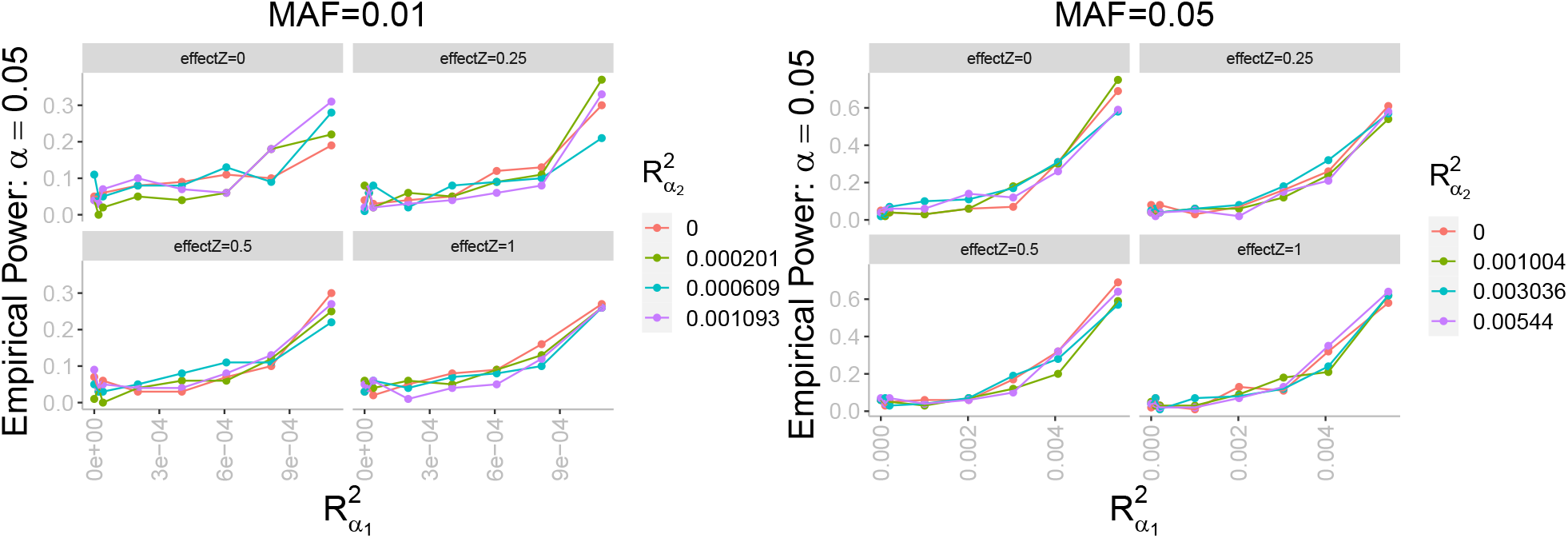
Power simulations: Scenario 2. Empirical power for GAMuT-Dissect as a function of the *R*^2^ corresponding to the effect *α*_1_ under Scenario 2. Different colors for different *R*^2^ for the effect *α*_2_ (no major differences by color). Simulated datasets (100) assumed a window of 500 rare genetic variants with a given MAF (0.01 left and 0.05 right). Different panels correspond to different polygenic effects (*Z*). Bigger polygenic effects of *Z* correspond to higher correlation between the phenotypes.

**Figure 5:**
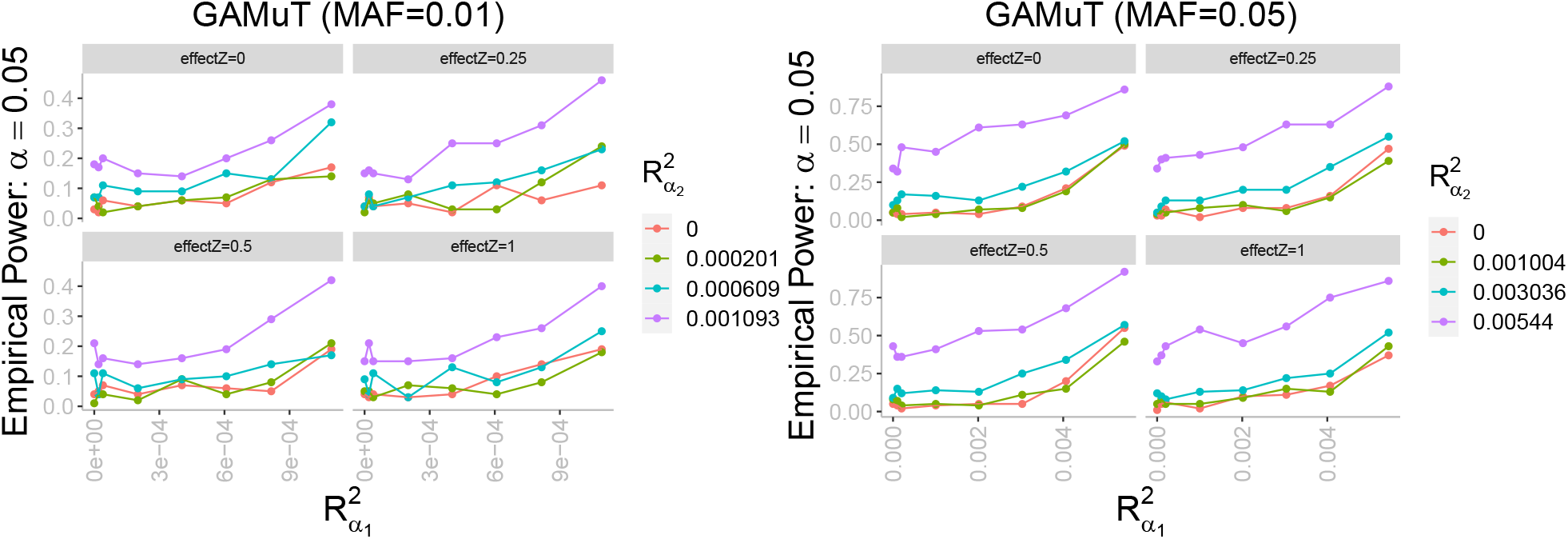
Power simulations: standard GAMuT association. Empirical power for standard GAMuT as a function of the *R*^2^ corresponding to the effect *α*_1_ under Scenario 2. Different colors for different *R*^2^ for the effect *α*_2_. Simulated datasets (100) assumed a window of genetic variants with a given MAF (0.01 left and 0.05 right). Different panels correspond to different polygenic effects (*Z*). Bigger polygenic effects of *Z* correspond to higher correlation between the phenotypes.

### 3.3 Power simulations (Scenario 3)

Here, we show that GAMuT-Dissect has power to detect Scenario 3 (see Figure 2 right) where the genetic variants have direct effects only on outcome variables after adjustment for polygenic effects *Z*. We present power plots as a function of *R*^2^ corresponding to *α*_1_, because this is the effect between *G* and the outcome. Figure 6 shows that GAMuT-Dissect has increasing power as the effect *α*_1_ increases. We also show that the parameters chosen for the effect size (*α*_1_) produce datasets with enough power to detect omnibus association with *G* (standard GAMuT in Figure 7).

**Figure 6:**
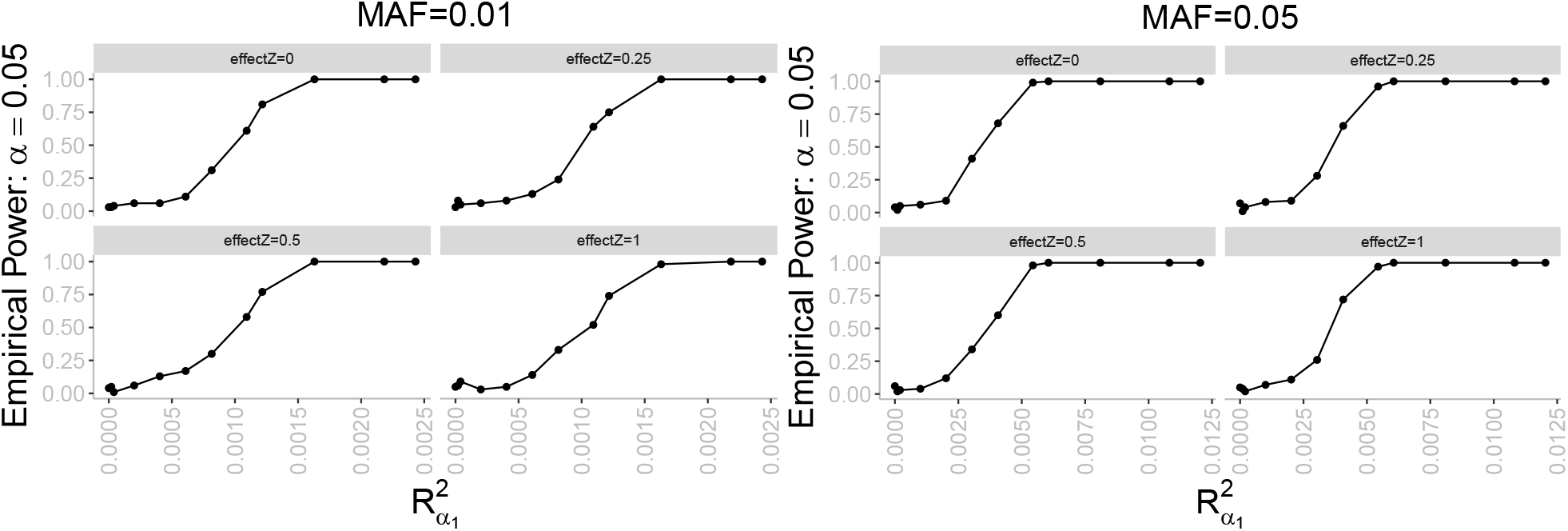
Power simulations: Scenario 3. Empirical power for GAMuT-Dissect as a function of the *R*^2^ corresponding to the effect *α*_1_ in the case of Scenario 3. Simulated datasets (100) assumed a window of genetic variants with a given MAF (0.01 left and 0.05 right). Different panels correspond to different polygenic effects (*Z*). Bigger polygenic effects of *Z* correspond to higher correlation between the phenotypes.

**Figure 7:**
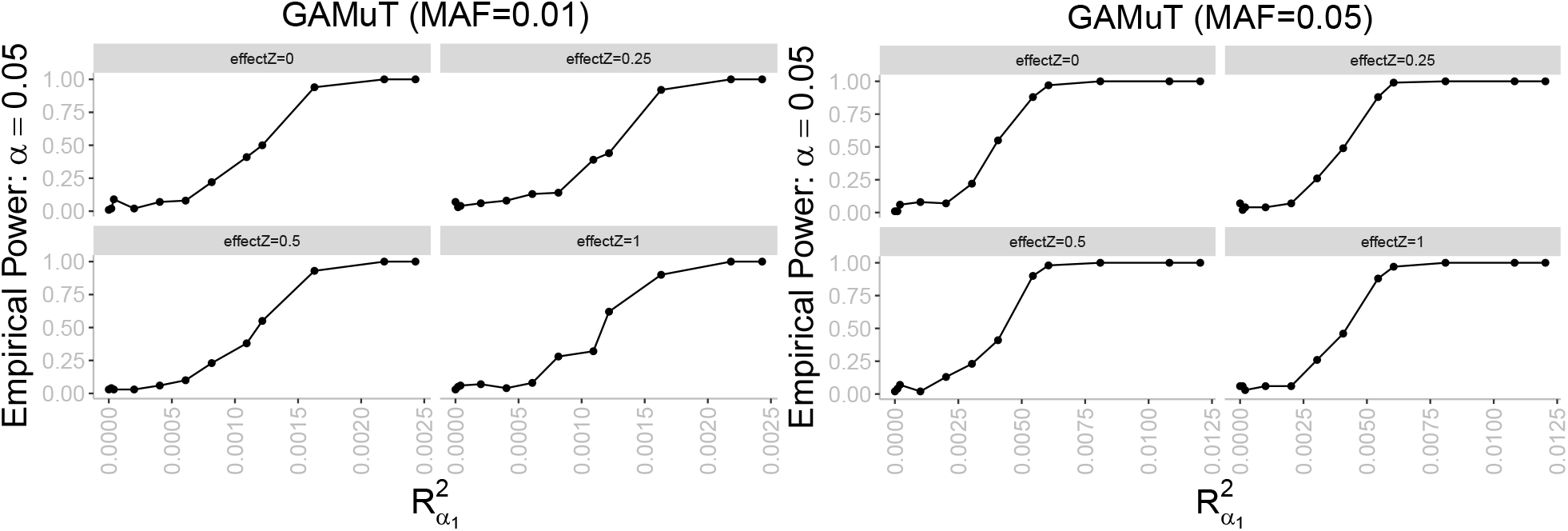
Power simulations: standard GAMuT association. Empirical power for standard GAMuT as a function of the *R*^2^ corresponding to the effect *α*_1_ in the case of Scenario 3. Simulated datasets (100) assumed a window of genetic variants with a given MAF (0.01 left and 0.05 right). Different panels correspond to different polygenic effects (*Z*). Bigger polygenic effects of *Z* correspond to higher correlation between the phenotypes.

### 3.4 Analysis of the Grady Trauma Project

With the GTP dataset, we applied GAMuT-Dissect by sequentially setting each of the 21 items composing the Beck Depression Inventory as a potential mediator between the variants in gene ZHX2 and the remaining 20 items in order to test whether particular BDI items were driving the association with ZHX2. Just as in Holleman et al. (2019), we controlled for gender, age, and ancestry, and used the estimated log odds ratios from external GWAS from the Psychiatric Genomics Consortium (Consortium et al., 2012; Group et al., 2011; Consortium et al., 2014) as the variants’ weights in GAMuT. We found that ZHX2 variants appeared to have direct effects on 3 items if we use the standard significance threshold of 0.05 (red line in Figure 8): self-dislike, worthlessness, irritability. This result means that there is evidence (at the 5% significance level) that the ZHX2 gene perhaps is directly associated with these symptoms (self-dislike, worthlessness, irritability), while having only weak or no direct associations with the other 18. By applying a Bonferroni multiple testing correction, we identify 8 mediator candidates (blue line in Figure 8): pessimism, self-dislike, self-criticalness, agitation, indecisiveness, worthlessness, irritability, concentration.

**Figure 8:**
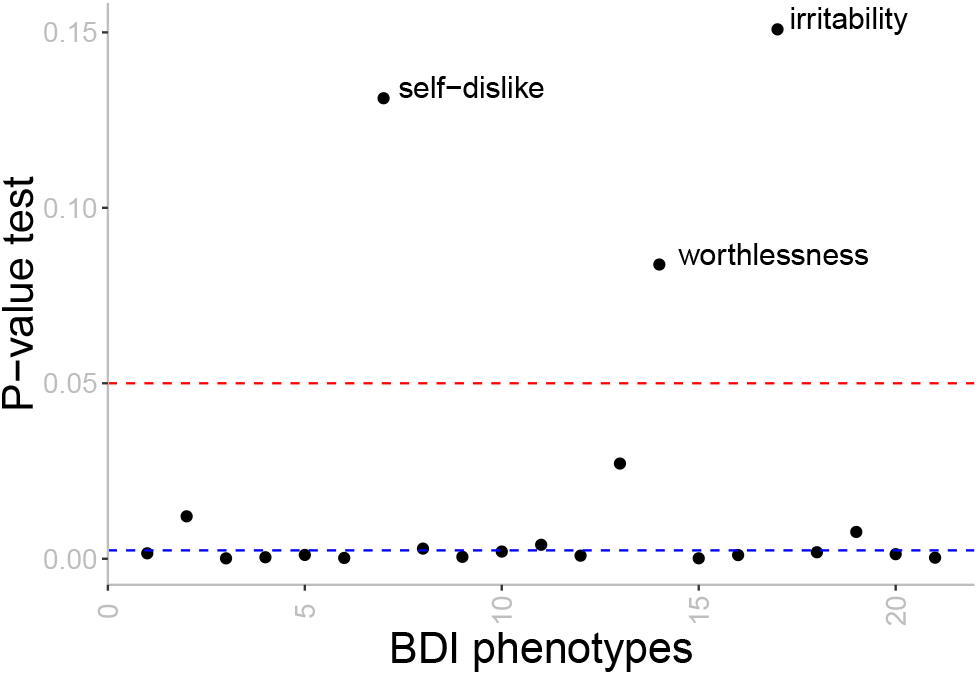
GTP tests. P-values of GAMuT-Dissect using each phenotype of GTP as a potential mediator one at a time between the ZHX2 gene and the other phenotypes in the BDI questionnaire (after adjusting for covariates). The null hypothesis of Scenario 1 is not rejected in three cases (significance level 5% (red line) and Bonferroni-corrected 0.23% (blue line)).

We next assessed whether the group of 8 items identified in our sequential analysis fully explains the association between ZHX2 and the complete set of 21 items. When we perform this new GAMuT-Dissect test of whether the set of all 8 phenotypes are directly associated with gene ZHX2, and the gene is not associated with the other 13 phenotypes, we again fail to reject the null hypothesis (Scenario 1) at the standard threshold level of 0.05 with p-value 0.1815221. Thus, there is evidence that these 8 phenotypes fully account for the association we originally observed in our omnibus GAMuT test between the ZHX2 gene and the 21 items that compose the BDI. This provides extra information that the standard GAMuT (or another standard pleitropy test) is unable to elucidate. While standard pleitropy tests are able to identify potential associations of one gene (ZHX2) with a collection of phenotypes (in this case, the 21 BDI symptoms), these tests cannot tease apart which phenotypes are directly associated with the gene and are driving the original multivariate signal. With our proposed GAMuT-Dissect, we are able to distinguish that there is evidence for a direct effect between gene ZHX2 and 8 symptoms (pessimism, self-dislike, self-criticalness, agitation, indecisiveness, worthlessness, irritability, concentration), while the remaining 13 symptoms do not appear directly influenced by the gene. We conducted a standard GAMuT between ZHX2 and these remaining 13 symptoms, which yielded a suggestive p-value of 5.5e-4. This result indicates that ZHX2 has potential indirect effects on at least a subset of the 13 symptoms mediated by the 8 directly associated symptoms identified by GAMuT-Dissect. We further tested if each of the 8 phenotypes was a mediator of the other 7. We identified 3 symptoms that fully mediate the relationship with the ZHX2 gene of the remaining 7 symptoms for a significance level of 0.625% (Bonferroni correction): self-dislike, worthlessness and irritability. Next, we tested whether those 3 symptoms fully mediated the relationship with ZHX2 of the remaining 5 and we were unable to reject full mediation with a p-value of 0.8917.

A followup study could focus on these 3 symptoms, and a more detailed phenotyping endevour, which could potentially help identify a more fine-grained collection of symptoms concentrated on self-dislike, worthlessness, irritability that are directly associated with the ZHX2 gene.

## 4 Discussion

A variety of genetic studies employ multivariate tests of multiple correlated phenotypes to exploit likely pleiotropy to improve power. In this work, we focus on the topic of follow-up genetic study of a significant omnibus test of cross-phenotype association. We propose a novel method built on the GAMuT framework for cross-phenotype testing of a gene set using kernel distance-covariance techniques. Our new kernel method, called GAMuT-Dissect, teases apart the significant signal from an omnibus test in order to determine the subset of phenotypes where the genetic variant (or set of variants) has a direct effect. Such information can yield improved biological insight of the traits under study.

Unlike existing mediation techniques that can tease apart direct and indirect genetic effects, GAMuT-Dissect can handle both high-dimensional phenotypic and genotypic data. Our method can also handle both categorical and continuous data and further adjusts for covariates. Furthermore, since GAMuT derives analytic P-values from Davies’ exact method, our proposed methodology is computationally efficient and applicable on a genome-wide scale. We simulated phenotypic data under a variety of direct and indirect genetic effects and showed our approach had good behavior across a wide range of models. We further illustrated our method using GWAS data from the Grady Trauma Project and showed that an existing signal between genetic variants in the ZHX2 gene and 21 items within the Beck Depression Inventory (Holleman et al., 2019) appears to be due to a direct effect of these variants on only 3 of these items.

We conclude with some comments on the characteristics of our method.

### Adjustment due to shared measured or unmeasured effects

First, our method requires adjustment for any shared effects (due to a common cause) among phenotypes that are independent of the genetic variants of interest. Failure to adjust for these shared effects (or similar covariates) can lead to misleading inference. If these shared effects are measured, we can regress such effects out prior to analysis. If the shared effects are random (i.e. modeling the polygenic correlations among phenotypes using genomewide correlation estimates from Bulik-Sullivan et al. 2015), we can decorrelate the phenotypes prior to analysis using the technique of (Zhou and Stephens, 2014) that eigendecomposes the resulting correlation/covariance matrix and multiplies the phenotypes by the eigenvector matrix.

### Interpreting the omnibus association

As noted when describing our method, GAMuT-Dissect tests the null that the omnibus association is completely explained by the subset of phenotypes modeled as the intermediate variables. Thus, when applying our method in general, special care should be used in interpreting the findings when rejecting or failing to reject the null, especially in relation to standard mediation techniques.

### Potential collider bias

Regarding the fact that GAMuT-Dissect teases apart the direct effects of *G* by conditioning on intermediate variables *P*_2_ (see Figure 1), we urge caution in interpretation of GAMuT-Dissect if there is reason to believe the outcome variables *P*_1_ causes *P*_2_ since the conditioning performed as part of our framework would then lead to collider bias. Thus, the underlying causal structure should be thought about before conditioning on a certain intermediate, in order to avoid a biased result.

### Greedy sequential approach to identify the mediator set

In this work, we assumed that the subsets *P*_1_ and *P*_2_ (mediator) are known in advance. This is likely not the case in some real-life applications. When the set of mediators is not known, one alternative is to devise a sequential approach in which first each individual phenotype is tested as a potential mediator (with the remaining *L* – 1 in *P*_1_). Then, the set of potential mediators is created by including those phenotypes identified as mediators by the GAMuT-Dissect test. In a second stage, pairs of mediators are tested together in *P*_2_ (with the remaining *L* – 2 phenotypes in *P*_1_). This same procedure is followed until the largest possible set of mediators is identified by GAMuT-Dissect. More is need to be studied of this greedy approach and will be left as future work. In particular, it is necessary to explore the control of type I error under this sequential approach that includes multiple (correlated) tests.

## Acknowledgements

This work was supported by NIH grants GM117946, HG007508, MH071537, and AR060893. We appreciate the technical support of all of the staff and volunteers of the Grady Trauma Project. Most importantly, we are extremely indebted to and appreciative of the time and effort given from all of the participants of the Grady Trauma Project.

## Web Resources

The URLs for software: https://github.com/crsl4/gamut-dissect and https://github.com/epstein-software.

## Data Availability

The Grady Trauma Project dataset is available from KJR on reasonable request.

## Supplementary material

### Type I Error Simulations: Single genetic variant

Figure 9 shows the quantile-quantile (QQ) plots of 100 null simulations of no association (*α*_1_ = *α*_2_ = 0 in simulation scenario figure in main text) for the standard GAMuT test (Broadaway et al., 2016) and for the mediation GAMuT test. Both methods properly control the type I error under the scenario of no association.

**Figure 9:**
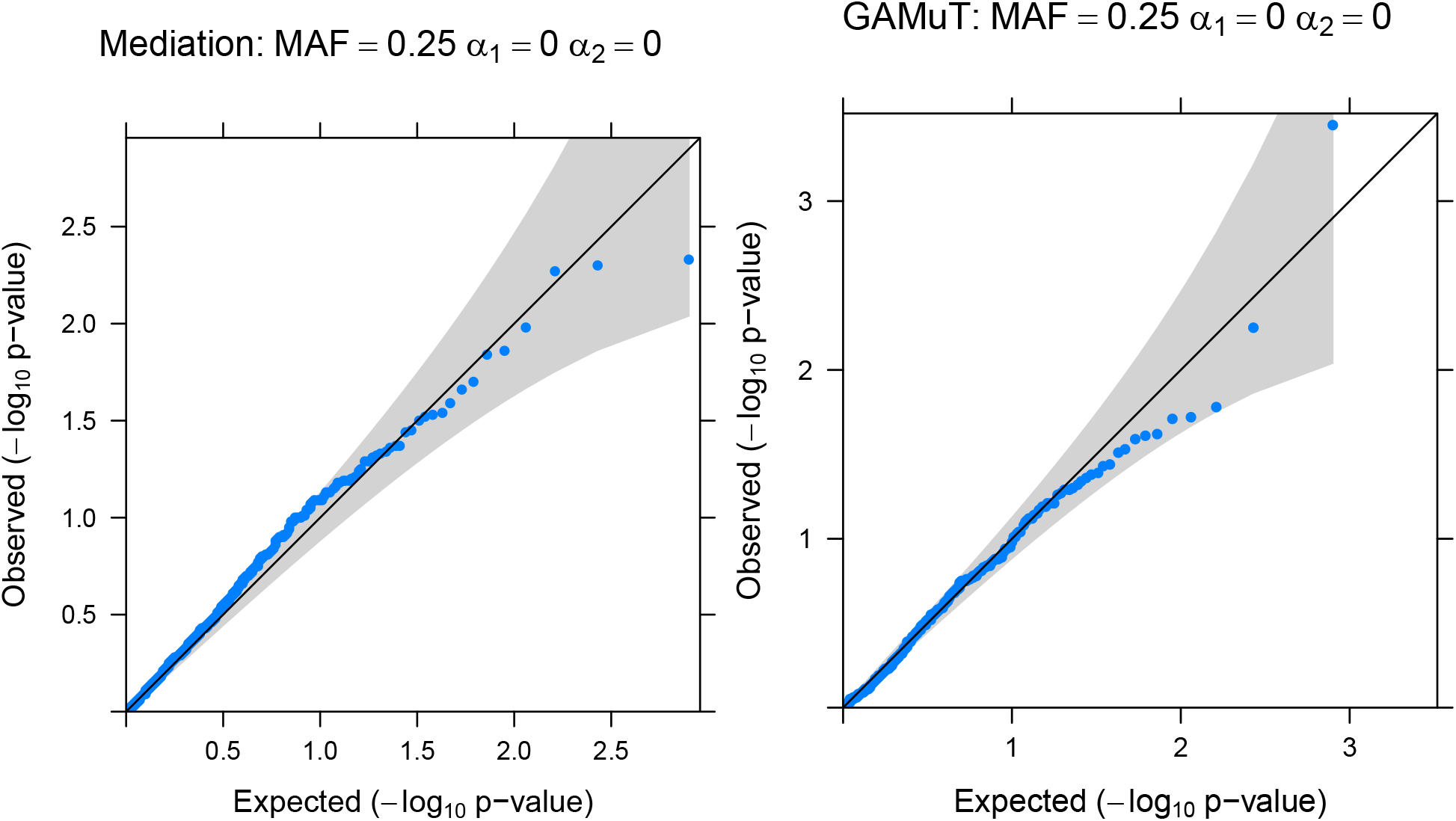
No association with single genetic variant. Q-Q plots of p-values for null simulations of no association (*α*_1_ = *α*_2_ = 0) for standard GAMuT and mediation GAMuT here proposed. Simulated datasets (100) assumed a single genetic variant with a given MAF.

Figure 10 shows the quantile-quantile (QQ) plots of 100 null simulations of full mediation (*α*_1_ =0 in simulation scenario figure in the main text) for the mediation GAMuT test here proposed when we adjust by the the polygenic effect (*Z*) on all the phenotypes (as described in the algorithm in the main text), and when we do not adjust by these effects. If we do not adjust by *Z*, there is inflation in the p-values (figure 10 left) when the genetic variant is of common variation (*MAF* = 0.25).

**Figure 10:**
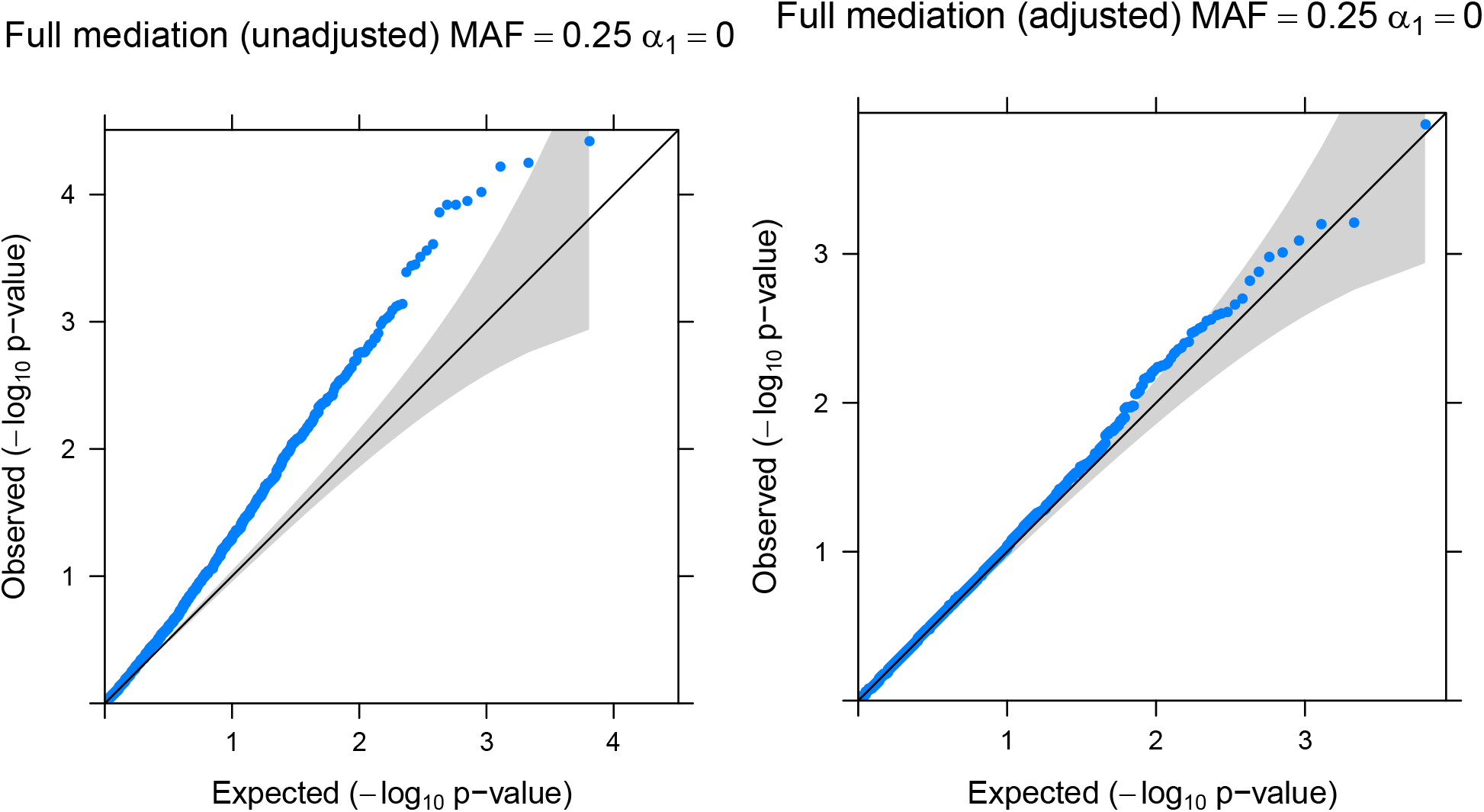
Full mediation with single genetic variant. Q-Q plots of p-values for null simulations of full mediation (*α*_1_ = 0) for mediation GAMuT here proposed. Simulated datasets (100) assumed a single genetic variant with a given MAF (0.01 or. Plots on the right are not adjusted by the polygenic effect (*Z*), and thus show inflation in p-values compared to the adjusted plots (left).

### Power Simulations of partial mediation: Single genetic variant

Here, we show that the mediation GAMuT has power to detect partial mediation. We present power plots as a function of *R*^2^ corresponding to *α*_1_, because this effect is the one driving the partial mediation, with different lines for *R*^2^ corresponding to *α*_2_. Figure 11 shows that the mediation GAMuT has increasing power as the effect *α*_1_ increases, regardless of the value of *α*_2_. We also show that the parameters chosen for the effect sizes (*α*_1_*, α*_2_) produce datasets with enough power to detect association with *G* (standard GAMuT in figure 12).

**Figure 11:**
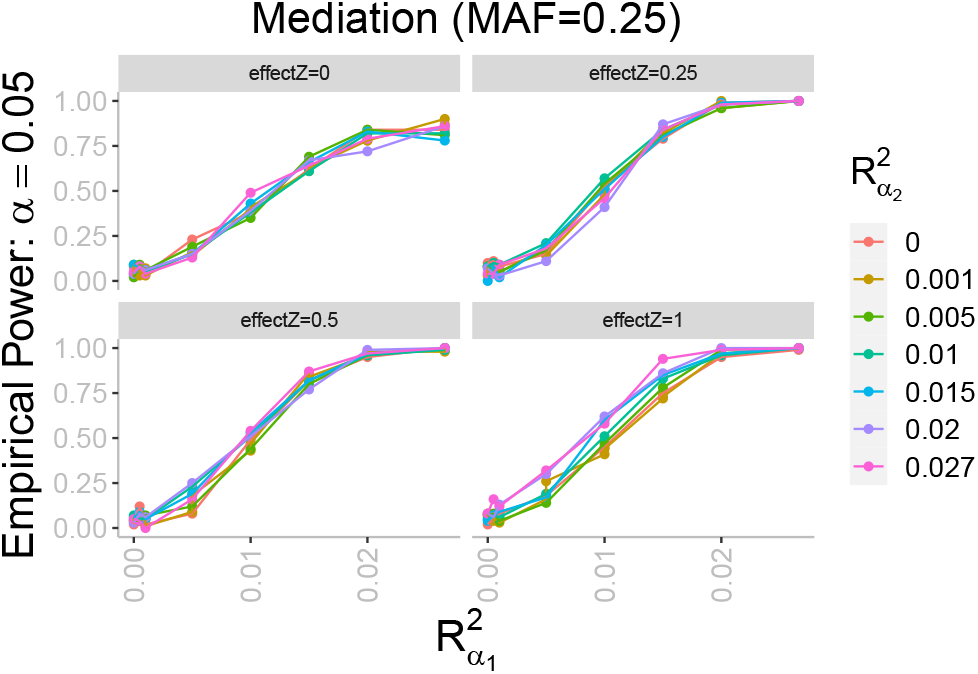
Partial mediation with single genetic variant. Empirical power for mediation GAMuT as a function of *R*^2^ corresponding to the effect *α*_1_ under partial mediation. Different colors for different *R*^2^ for the effect *α*_2_ (no major differences by color). Simulated datasets (100) assumed a single genetic variant with a given MAF. Different panels correspond to different polygenic effects (*Z*). Bigger polygenic effects of *Z* correspond to higher correlation among the phenotypes.

**Figure 12:**
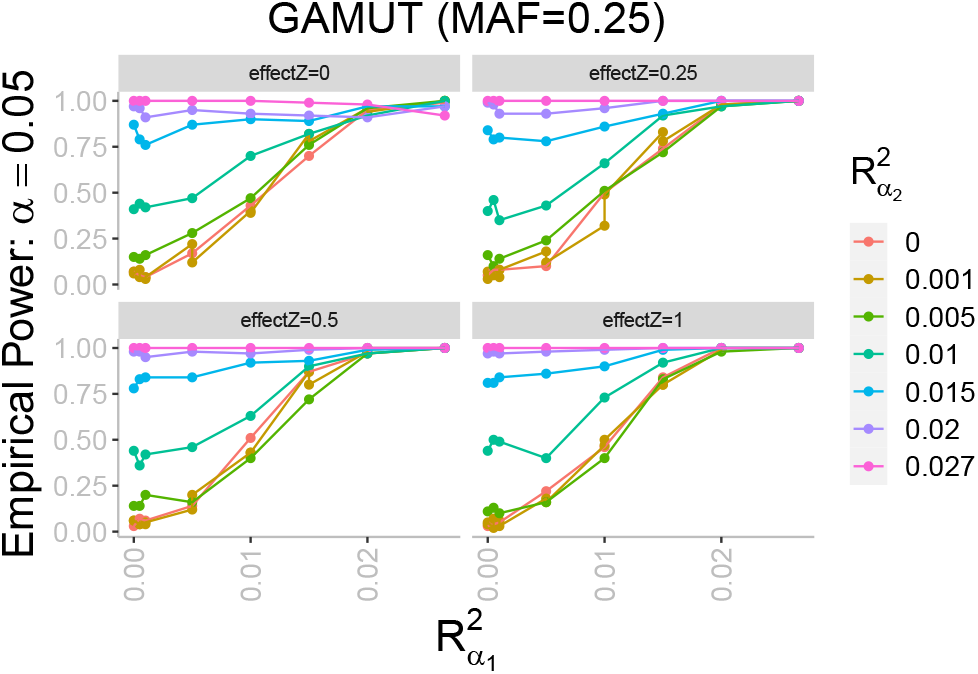
Association with single genetic variant. Empirical power for standard GAMuT as a function of *R*^2^ corresponding to the effect *α*_1_ under partial mediation. Different colors for different *R*^2^ for the effect *α*_2_ (which contributes to the association). Simulated datasets (100) assumed a single genetic variant with a given MAF. Different panels correspond to different polygenic effects (*Z*). Bigger polygenic effects of *Z* correspond to higher correlation between the phenotypes.

### Power Simulations of no mediation: Single genetic variant

Here, we show that the mediation GAMuT does not have power to distinguish no mediation from the full mediation (null hypothesis) for the case of correlated phenotypes when we estimate the polygenic effect *Z* with *PC*1. We present power plots as a function of *R*^2^ corresponding to *α*_1_ (which we assume to be the same effect for both *Y*_1_ and *Y*_2_ in this scenario for simplicity).

In this setup, the mediator set [*Y*_3_, *Y*_4_] (simulation scenario figure in main text) is not associated with *G*. When the phenotypes are uncorrelated (effect of *Z* is zero in figure 13), the empirical power has a peak at a certain effect size *α*_1_. We suspect that as the effect size (*α*_1_) increases, the PC represents this signal instead of the effect from *Z*. Thus, when we residualize by the PCs, we lose the association signal. When the phenotypes are correlated (effect of *Z* greater than zero), suddenly the empirical power is close to the significance level, hence, the test does not have power to detect between the alternative hypothesis of no mediation and the null hypothesis of full mediation. This situation is not created by lack of signal of an association between *G* and the set of all phenotypes ([*Y*_1_, *Y*_2_, *Y*_3_, *Y*_4_]) as shown in figure 14.

**Figure 13:**
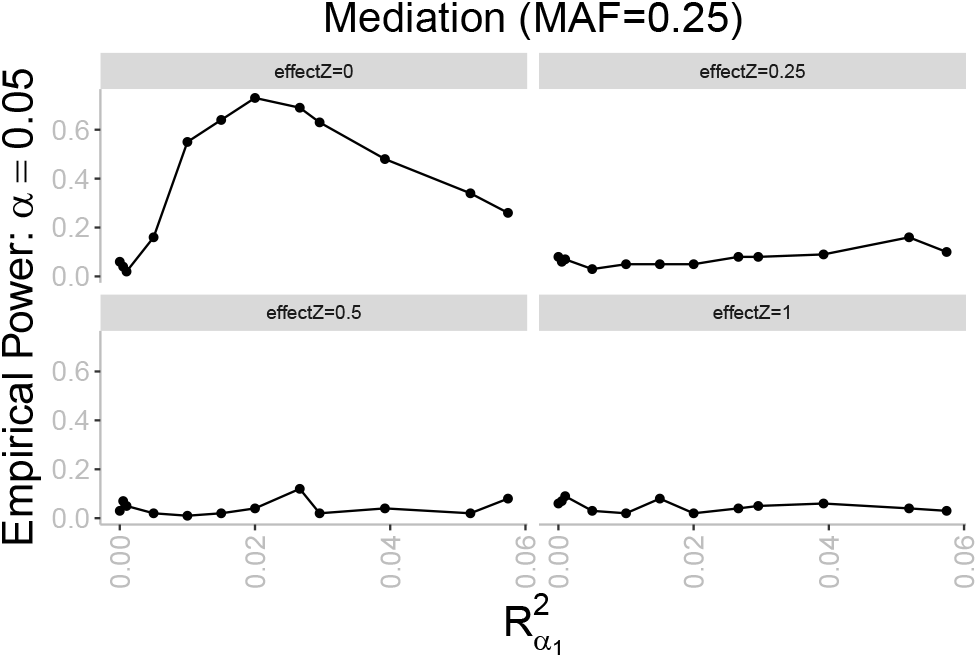
No mediation with single genetic variant. Empirical power for mediation GAMuT as a function of the *R*^2^ corresponding to the effect *α*_1_ in the case of no mediation. Simulated datasets (100) assumed a single genetic variant with a given MAF. Different panels correspond to different polygenic effects (*Z*). Bigger polygenic effects of *Z* correspond to higher correlation between the phenotypes.

**Figure 14:**
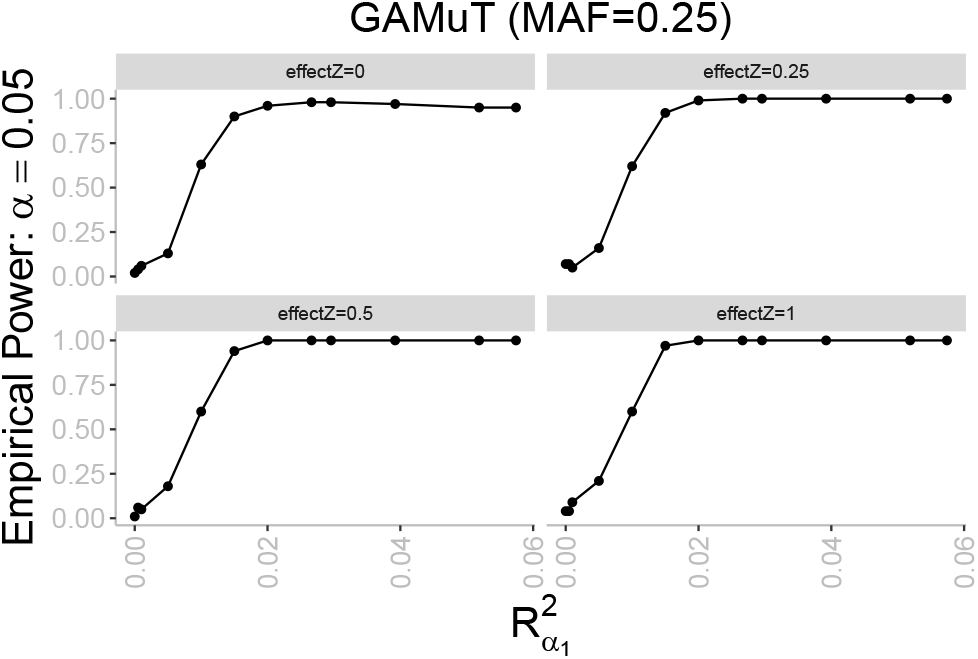
No mediation with single genetic variant. Empirical power for standard GAMuT as a function of the *R*^2^ corresponding to the effect *α*_1_ in the case of no mediation. Simulated datasets (100) assumed a single genetic variant with a given MAF. Different panels correspond to different polygenic effects (*Z*). Bigger polygenic effects of *Z* correspond to higher correlation between the phenotypes.

## References

Bradley, R. G., Binder, E. B., Epstein, M. P., Tang, Y., Nair, H. P., Liu, W., Gillespie, C. F., Berg, T., Evces, M., Newport, D. J., Stowe, Z. N., Heim, C. M., Nemeroff, C. B., Schwartz, A., Cubells, J. F., and Ressler, K. J. (2008). Influence of Child Abuse on Adult Depression: Moderation by the Corticotropin-Releasing Hormone Receptor Gene. JAMA Psychiatry, 65(2):190–200.

Broadaway, K. A., Cutler, D. J., Duncan, R., Moore, J. L., Ware, E. B., Jhun, M. A., Bielak, L. F., Zhao, W., Smith, J. A., Peyser, P. A., Kardia, S. L. R., Ghosh, D., and Epstein, M. P. (2016). A statistical approach for testing cross-phenotype effects of rare variants. American Journal of Human Genetics, 98(3):525–540.

Chung, D., Yang, C., Li, C., Gelernter, J., and Zhao, H. (2014). GPA: A Statistical Approach to Prioritizing GWAS Results by Integrating Pleiotropy and Annotation. PLOS Genetics, 10(11):e1004787.

Consortium, M. D. D. W. G. o. t. P. G., Ripke, S., Wray, N. R., Lewis, C. M., Hamilton, S. P., Weissman, M. M., Breen, G., Byrne, E. M., Blackwood, D. H. R., Boomsma, D. I., Cichon, S., Heath, A. C., Holsboer, F., Lucae, S., Madden, P. A. F., Martin, N. G., McGuffin, P., Muglia, P., Noethen, M. M., Penninx, B.P., Pergadia, M. L., Potash, J. B., Rietschel, M., Lin, D., Müller-Myhsok, B., Shi, J., Steinberg, S., Grabe, H. J., Lichtenstein, P., Magnusson, P., Perlis, R. H., Preisig, M., Smoller, J. W., Stefansson, K., Uher, R., Kutalik, Z., Tansey, K. E., Teumer, A., Viktorin, A., Barnes, M. R., Bettecken, T., Binder, E. B., Breuer, R., Castro, V. M., Churchill, S. E., Coryell, W. H., Craddock, N., Craig, I. W., Czamara, D., De Geus, E. J., Degenhardt, F., Farmer, A. E., Fava, M., Frank, J., Gainer, V. S., Gallagher, P. J., Gordon, S. D., Goryachev, S., Gross, M., Guipponi, M., Henders, A. K., Herms, S., Hickie, I. B., Hoefels, S., Hoogendijk, W., Hottenga, J. J., Iosifescu, D. V., Ising, M., Jones, I., Jones, L., Jung-Ying, T., Knowles, J. A., Kohane, I. S., Kohli, M. A., Korszun, A., Landen, M., Lawson, W. B., Lewis, G., MacIntyre, D., Maier, W., Mattheisen, M., McGrath, P. J., McIntosh, A., McLean, A., Middeldorp, C. M., Middleton, L., Montgomery, G. M., Murphy, S. N., Nauck, M., Nolen, W. A., Nyholt, D. R., O’Donovan, M., Oskarsson, H., Pedersen, N., Scheftner, W. A., Schulz, A., Schulze, T. G., Shyn, S. I., Sigurdsson, E., Slager, S. L., Smit, J. H., Stefansson, H., Steffens, M., Thorgeirsson, T., Tozzi, F., Treutlein, J., Uhr, M., van den Oord, E.J. C. G., Van Grootheest, G., Völzke, H., Weilburg, J. B., Willemsen, G., Zitman, F. G., Neale, B., Daly, M., Levinson, D. F., and Sullivan, P. F. (2012). A mega-analysis of genome-wide association studies for major depressive disorder. Molecular Psychiatry, 18:497.

Consortium, S. W. G. o. t. P. G., Ripke, S., Neale, B. M., Corvin, A., Walters, J. T. R., Farh, K.-H., Holmans, P. A., Lee, P., Bulik-Sullivan, B., Collier, D. A., Huang, H., Pers, T. H., Agartz, I., Agerbo, E., Albus, M., Alexander, M., Amin, F., Bacanu, S. A., Begemann, M., Belliveau Jr, R. A., Bene, J., Bergen, S. E., Bevilacqua, E., Bigdeli, T. B., Black, D. W., Bruggeman, R., Buccola, N. G., Buckner, R. L., Byerley, W., Cahn, W., Cai, G., Campion, D., Cantor, R. M., Carr, V. J., Carrera, N., Catts, S. V., Chambert, K. D., Chan, R. C. K., Chen, R. Y. L., Chen, E. Y. H., Cheng, W., Cheung, E. F. C., Ann Chong, S., Robert Cloninger, C., Cohen, D., Cohen, N., Cormican, P., Craddock, N., Crowley, J. J., Curtis, D., Davidson, M., Davis, K. L., Degenhardt, F., Del Favero, J., Demontis, D., Dikeos, D., Dinan, T., Djurovic, S., Donohoe, G., Drapeau, E., Duan, J., Dudbridge, F., Durmishi, N., Eichhammer, P., Eriksson, J., Escott-Price, V., Essioux, L., Fanous, A. H., Farrell, M. S., Frank, J., Franke, L., Freedman, R., Freimer, N. B., Friedl, M., Friedman, J. I., Fromer, M., Genovese, G., Georgieva, L., Giegling, I., Giusti-Rodríguez, P., Godard, S., Goldstein, J. I., Golimbet, V., Gopal, S., Gratten, J., de Haan, L., Hammer, C., Hamshere, M. L., Hansen, M., Hansen, T., Haroutunian, V., Hartmann, A. M., Henskens, F. A., Herms, S., Hirschhorn, J. N., Hoffmann, P., Hofman, A., Hollegaard, M. V., Hougaard, D. M., Ikeda, M., Joa, I., Juliá, A., Kahn, R. S., Kalaydjieva, L., Karachanak-Yankova, S., Karjalainen, J., Kavanagh, D., Keller, M. C., Kennedy, J. L., Khrunin, A., Kim, Y., Klovins, J., Knowles, J. A., Konte, B., Kucinskas, V., Ausrele Kucinskiene, Z., Kuzelova-Ptackova, H., Kähler, A. K., Laurent, C., Lee Chee Keong, J., Hong Lee, S., Legge, S. E., Lerer, B., Li, M., Li, T., Liang, K.-Y., Lieberman, J., Limborska, S., Loughland, C. M., Lubinski, J., Lönnqvist, J., Macek Jr, M., Magnusson, P. K. E., Maher, B. S., Maier, W., Mallet, J., Marsal, S., Mattheisen, M., Mattingsdal, M., McCarley, R. W., McDonald, C., McIntosh, A. M., Meier, S., Meijer, C. J., Melegh, B., Melle, I., Mesholam-Gately, R. I., Metspalu, A., Michie, P. T., Milani, L., Milanova, V., Mokrab, Y., Morris, D. W., Mors, O., Murphy, K. C., Murray, R. M., Myin-Germeys, I., Müller-Myhsok, B., Nelis, M., Nenadic, I., Nertney, D. A., Nestadt, G., Nicodemus, K. K., Nikitina-Zake, L., Nisenbaum, L., Nordin, A., O’Callaghan, E., O’Dushlaine, C., O’Neill, F. A., Oh, S.-Y., Olincy, A., Olsen, L., Van Os, J., Pantelis, C., Papadimitriou, G. N., Papiol, S., Parkhomenko, E., Pato, M. T., Paunio, T., Pejovic-Milovancevic, M., Perkins, D. O., Pietiläinen, O., Pimm, J., Pocklington, A. J., Powell, J., Price, A., Pulver, A. E., Purcell, S. M., Quested, D., Rasmussen, H. B., Reichenberg, A., Reimers, M. A., Richards, A. L., Roffman, J. L., Roussos, P., Ruderfer, D. M., Salomaa, V., Sanders, A. R., Schall, U., Schubert, C. R., Schulze, T.G., Schwab, S. G., Scolnick, E. M., Scott, R. J., Seidman, L. J., Shi, J., Sigurdsson, E., Silagadze, T., Silverman, J. M., Sim, K., Slominsky, P., Smoller, J. W., So, H.-C., Spencer, C. A., Stahl, E. A., Stefansson, H., Steinberg, S., Stogmann, E., Straub, R. E., Strengman, E., Strohmaier, J., Scott Stroup, T., Subramaniam, M., Suvisaari, J., Svrakic, D. M., Szatkiewicz, J. P., Söderman, E., Thirumalai, S., Toncheva, D., Tosato, S., Veijola, J., Waddington, J., Walsh, D., Wang, D., Wang, Q., Webb, B. T., Weiser, M., Wildenauer, D. B., Williams, N. M., Williams, S., Witt, S. H., Wolen, A. R., Wong, E. H. M., Wormley, B. K., Simon Xi, H., Zai, C. C., Zheng, X., Zimprich, F., Wray, N. R., Stefansson, K., Visscher, P. M., Trust Case-Control Consortium, W., Adolfsson, R., Andreassen, O. A., Blackwood, D. H. R., Bramon, E., Buxbaum, J. D., Børglum, A. D., Cichon, S., Darvasi, A., Domenici, E., Ehrenreich, H., Esko, T., Gejman, P. V., Gill, M., Gurling, H., Hultman, C. M., Iwata, N., Jablensky, A. V., Jönsson, E. G., Kendler, K. S., Kirov, G., Knight, J., Lencz, T., Levinson, D. F., Li, Q. S., Liu, J., Malhotra, A. K., McCarroll, S. A., McQuillin, A., Moran, J. L., Mortensen, P. B., Mowry, B. J., Nöthen, M. M., Ophoff, R. A., Owen, M. J., Palotie, A., Pato, C. N., Petryshen, T. L., Posthuma, D., Rietschel, M., Riley, B. P., Rujescu, D., Sham, P. C., Sklar, P., St Clair, D., Weinberger, D. R., Wendland, J. R., Werge, T., Daly, M. J., Sullivan, P. F., and O’Donovan, M. C. (2014). Biological insights from 108 schizophrenia-associated genetic loci. Nature, 511:421.

Davies, R. B. (1980). Algorithm as 155: The distribution of a linear combination of *χ*^2^ random variables. Journal of the Royal Statistical Society. Series C (Applied Statistics), 29(3):323–333.

Gabrielsen, M. E., Romundstad, P., Langhammer, A., Krokan, H. E., and Skorpen, F. (2013). Association between a 15q25 gene variant, nicotine-related habits, lung cancer and COPD among 56,307 individuals from the HUNT study in Norway. European journal of human genetics: EJHG, 21(11):1293–1299.

Galesloot, T. E., van Steen, K., Kiemeney, L. A. L. M., Janss, L. L., and Vermeulen, S. H. (2014). A comparison of multivariate genome-wide association methods. PloS one, 9(4):e95923–e95923.

Gretton, A., Fukumizu, K., Teo, C., Song, L., Schölkopf, B., and Smola, A. (2008). A kernel statistical test of independence. In Advances in neural information processing systems 20, pages 585–592, Red Hook, NY, USA. Max-Planck-Gesellschaft, Curran.

Group, P. G. C. B. D. W., Sklar, P., Ripke, S., Scott, L. J., Andreassen, O. A., Cichon, S., Craddock, N., Edenberg, H. J., Nurnberger Jr, J. I., Rietschel, M., Blackwood, D., Corvin, A., Flickinger, M., Guan, W., Mattingsdal, M., McQuillin, A., Kwan, P., Wienker, T. F., Daly, M., Dudbridge, F., Holmans, P. A., Lin, D., Burmeister, M., Greenwood, T. A., Hamshere, M. L., Muglia, P., Smith, E. N., Zandi, P. P., Nievergelt, C. M., McKinney, R., Shilling, P. D., Schork, N. J., Bloss, C. S., Foroud, T., Koller, D. L., Gershon, E. S., Liu, C., Badner, J. A., Scheftner, W. A., Lawson, W. B., Nwulia, E. A., Hipolito, M., Coryell, W., Rice, J., Byerley, W., McMahon, F. J., Schulze, T. G., Berrettini, W., Lohoff, F. W., Potash, J. B., Mahon, P. B., McInnis, M. G., Zöllner, S., Zhang, P., Craig, D. W., Szelinger, S., Barrett, T. B., Breuer, R., Meier, S., Strohmaier, J., Witt, S. H., Tozzi, F., Farmer, A., McGuffin, P., Strauss, J., Xu, W., Kennedy, J. L., Vincent, J. B., Matthews, K., Day, R., Ferreira, M. A., O’Dushlaine, C., Perlis, R., Raychaudhuri, S., Ruderfer, D., Lee, P. H., Smoller, J. W., Li, J., Absher, D., Bunney, W. E., Barchas, J. D., Schatzberg, A. F., Jones, E. G., Meng, F., Thompson, R. C., Watson, S. J., Myers, R. M., Akil, H., Boehnke, M., Chambert, K., Moran, J., Scolnick, E., Djurovic, S., Melle, I., Morken, G., Gill, M., Morris, D., Quinn, E., Mühleisen, T. W., Degenhardt, F. A., Mattheisen, M., Schumacher, J., Maier, W., Steffens, M., Propping, P., Nöthen, M. M., Anjorin, A., Bass, N., Gurling, H., Kandaswamy, R., Lawrence, J., McGhee, K., McIntosh, A., McLean, A. W., Muir, W. J., Pickard, B. S., Breen, G., St. Clair, D., Caesar, S., Gordon-Smith, K., Jones, L., Fraser, C., Green, E. K., Grozeva, D., Jones, I. R., Kirov, G., Moskvina, V., Nikolov, I., O’Donovan, M. C., Owen, M. J., Collier, D. A., Elkin, A., Williamson, R., Young, A. H., Ferrier, I. N., Stefansson, K., Stefansson, H., Borgeirsson, B., Steinberg, S., Gustafsson, Ó., Bergen, S. E., Nimgaonkar, V., Hultman, C., Landén, M., Lichtenstein, P., Sullivan, P., Schalling, M., Osby, U., Backlund, L., Frisén, L., Langstrom, N., Jamain, S., Leboyer, M., Etain, B., Bellivier, F., Petursson, H., Sigurdsson, E., Müller-Mysok, B., Lucae, S., Schwarz, M., Fullerton, J. M., Schofield, P. R., Martin, N., Montgomery, G. W., Lathrop, M., Óskarsson, H., Bauer, M., Wright, A., Mitchell, P. B., Hautzinger, M., Reif, A., Kelsoe, J. R., and Purcell, S. M. (2011). Large-scale genome-wide association analysis of bipolar disorder identifies a new susceptibility locus near ODZ4. Nature Genetics, 43:977.

Holleman, A. M., Broadaway, K. A., Duncan, R., Todor, A., Almli, L. M., Bradley, B., Ressler, K. J., Ghosh, D., Mulle, J. G., and Epstein, M. P. (2019). Powerful and Efficient Strategies for Genetic Association Testing of Symptom and Questionnaire Data in Psychiatric Genetic Studies. Scientific Reports, 9(1):7523.

Hua, W.-Y. and Ghosh, D. (2015). Equivalence of kernel machine regression and kernel distance covariance for multidimensional phenotype association studies. Biometrics, 71(3):812–820.

Huang, Y.-T., Vanderweele, T. J., and Lin, X. (2014). Joint analysis of SNP and gene expression data in genetic association studies of complex diseases. The annals of applied statistics, 8(1):352–376.

Kang, H. M., Sul, J. H., Service, S. K., Zaitlen, N. A., Kong, S.-y., Freimer, N. B., Sabatti, C., and Eskin, E. (2010). Variance component model to account for sample structure in genome-wide association studies. Nature genetics, 42(4):348–354.

Ma, C., Boehnke, M., Lee, S., and Investigators, G. (2015). Evaluating the calibration and power of three gene-based association tests of rare variants for the x chromosome. Genetic epidemiology, 39(7):499–508.

Price, A. L., Patterson, N. J., Plenge, R. M., Weinblatt, M. E., Shadick, N. A., and Reich, D. (2006). Principal components analysis corrects for stratification in genome-wide association studies. Nature Genetics, 38:904.

Ressler, K. J., Mercer, K. B., Bradley, B., Jovanovic, T., Mahan, A., Kerley, K., Norrholm, S. D., Kilaru, V., Smith, A. K., Myers, A. J., Ramirez, M., Engel, A., Hammack, S. E., Toufexis, D., Braas, K. M., Binder, E. B., and May, V. (2011). Post-traumatic stress disorder is associated with PACAP and the PAC1 receptor. Nature, 470:492.

Schaid, D. J., Tong, X., Larrabee, B., Kennedy, R. B., Poland, G. A., and Sinnwell, J. P. (2016). Statistical methods for testing genetic pleiotropy. Genetics, 204(2):483–497.

Solovieff, N., Cotsapas, C., Lee, P. H., Purcell, S. M., and Smoller, J. W. (2013). Pleiotropy in complex traits: challenges and strategies. Nature reviews. Genetics, 14(7):483–495.

Stevens, J. R., Al Masud, A., and Suyundikov, A. (2017). A comparison of multiple testing adjustment methods with block-correlation positively-dependent tests. PLOS ONE, 12(4):1–12.

Székely, G. J. and Rizzo, M. L. (2009). Brownian distance covariance. Ann. Appl. Stat., 3(4):1236–1265.

Székely, G. J., Rizzo, M. L., and Bakirov, N. K. (2007). Measuring and testing dependence by correlation of distances. Ann. Statist., 35(6):2769–2794.

VanderWeele, T. J., Asomaning, K., Tchetgen Tchetgen, E. J., Han, Y., Spitz, M. R., Shete, S., Wu, X., Gaborieau, V., Wang, Y., McLaughlin, J., Hung, R. J., Brennan, P., Amos, C. I., Christiani, D. C., and Lin, X. (2012). Genetic variants on 15q25.1, smoking, and lung cancer: an assessment of mediation and interaction. American journal of epidemiology, 175(10):1013–1020.

Walker, S. and Scherer, S. W. (2013). Identification of candidate intergenic risk loci in autism spectrum disorder. BMC Genomics, 14(1):499.

Wei, C. and Lu, Q. (2017). A generalized association test based on u statistics. Bioinformatics (Oxford, England), 33(13):1963–1971.

Zhou, X. and Stephens, M. (2014). Efficient multivariate linear mixed model algorithms for genome-wide association studies. Nature methods, 11(4):407–409.

